# RNA-triggered Cas12a2 antibacterial activity and resistance plasmid degradation

**DOI:** 10.64898/2026.11.03.763947

**Authors:** Yang Bai, Yifei Song, Wenjie Gu, Haiping Wu, Yunlong Liu

## Abstract

Antimicrobial resistance (AMR) necessitates programmable antibacterial strategies with novel mechanisms. Here, we evaluated an inducible intracellular CRISPR/Cas12a2 system in engineered *Escherichia coli* targeting a chloramphenicol-resistance (Chl) transcript. Cas12a2 activation significantly delayed bacterial growth and reduced recoverable colony-forming units (CFU) by 3-log_10_ *in vitro*, and reduced bacterial burdens in a mouse skin wound model. Transcriptomic profiling revealed upregulation of *recA*, a key mediator of the SOS response. We therefore designed a Chl-*recA* dual-targeting construct, which produced a greater growth delay than either single-targeting counterpart. The optimized dual-targeting system led to an approximately 4-log_10_ reduction in bacterial burden *in vivo*. Furthermore, exploratory qPCR of culture supernatants suggested a treatment-associated decrease in amplifiable plasmid DNA. Overall, this study provides proof-of-concept for resistance-transcript-guided Cas12a2 antibacterial activity and highlights the potential of co-targeting stress-associated transcripts to enhance programmable antimicrobials.

## Introduction

Antimicrobial resistance (AMR) threatens the effective treatment of bacterial infections and motivates the development of antibacterial strategies with mechanisms distinct from those of conventional antibiotics ^1, 2^. CRISPR-Cas systems offer a programmable approach by linking the recognition of defined nucleic acid sequences to bacterial growth inhibition or killing ^3-5^. Resistance-associated sequences are particularly relevant recognition targets because they can provide a molecular basis for distinguishing bacteria carrying specific resistance determinant.

Different CRISPR effectors connect sequence recognition to distinct forms of nucleic acid damage. DNA-targeting effectors can induce chromosomal cleavage or promote the loss of targeted plasmids, whereas RNA-targeting effectors can couple transcript recognition to collateral RNA degradation ^6-8^. Cas12a2 provides another route: recognition of a cognate RNA target activates collateral cleavage of RNA and DNA substrates ^9, 10^. This biochemical activity creates the possibility of using a resistance-gene transcript as a recognition input for antibacterial effects extending beyond cleavage of the initiating RNA. Bacterial responses to nucleic acid damage may also provide opportunities for additional recognition inputs. The SOS response coordinates changes in gene expression associated with DNA damage, repair and stress adaptation ^11, 12^. Although these responses generally support bacterial adaptation to damage, the resulting transcripts could potentially serve as targets for an RNA-triggered nuclease. We therefore considered whether combining recognition of a resistance-associated transcript with recognition of *recA* mRNA could improve Cas12a2 antibacterial activity. In this design, the resistance transcript provides one recognition input, while a stress-associated transcript provides a candidate second input.

Here, we investigate this design in an engineered *Escherichia coli* BL21(DE3) pLysS model carrying a chloramphenicol-resistance determinant (**Figure 1**). We evaluate inducible intracellular expression of Cas12a2 with a crRNA directed against the chloramphenicol-resistance transcript, using optical-density measurements and colony counts to distinguish changes in bacterial growth from changes in recoverable viable bacteria. Then, we use transcriptome profiling and reverse-transcription quantitative PCR to characterize SOS-associated transcriptional changes following Cas12a2 activation and to assess *recA* as a candidate secondary target. These observations motivate the comparison of dual-targeting constructs incorporating resistance-transcript and *recA* recognition, followed by evaluation of a selected construct in the wound model. Finally, we explored temporal changes in resistance-plasmid-associated DNA in culture supernatants during treatment.

**Figure 1.**
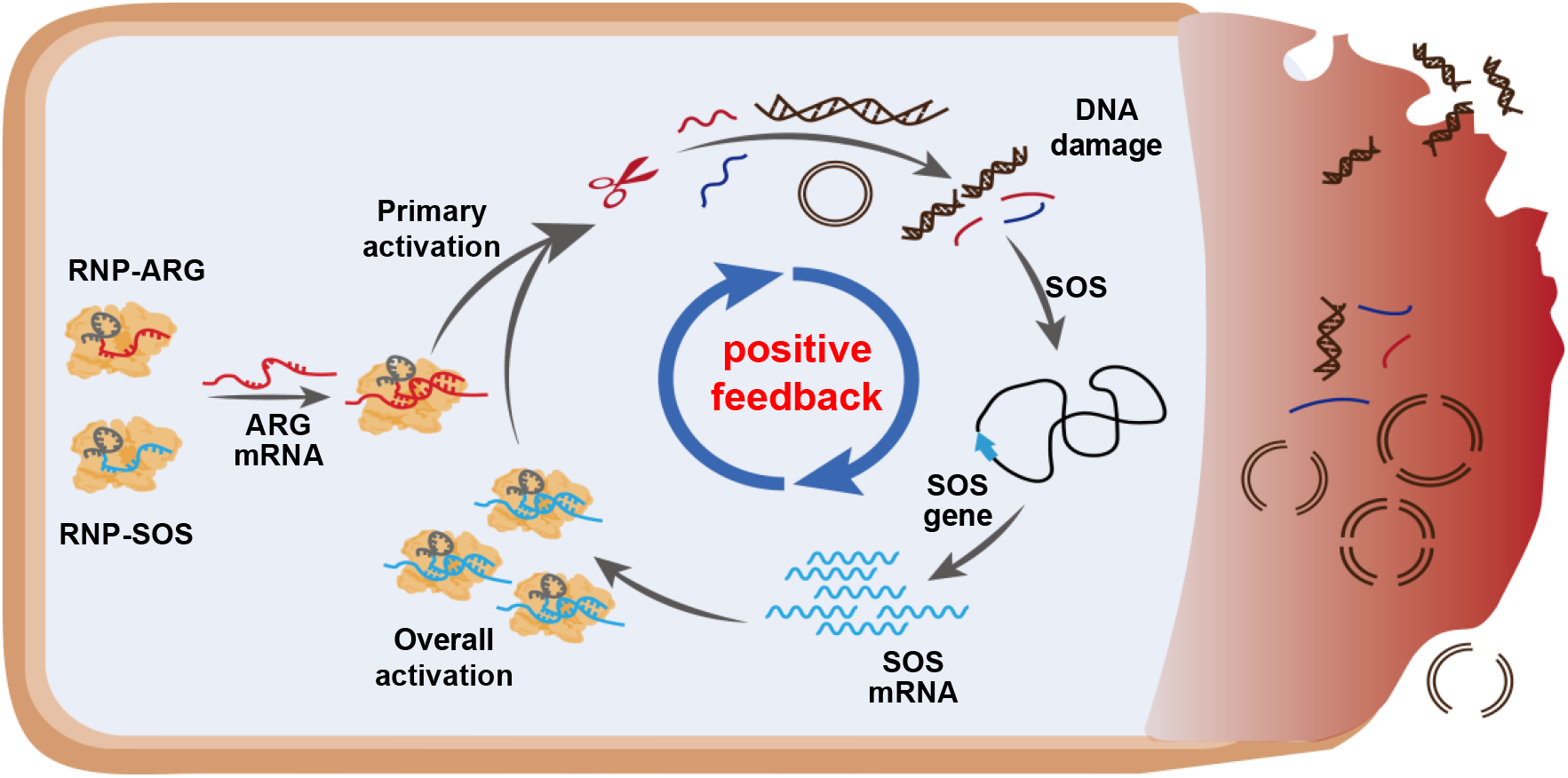
Overall strategy of ARG triggered, SOS-amplified antibacterial enhancement.

## Results and discussion

### Intracellular Cas12a2 targeting affects bacterial growth and viability in culture and bacterial burden in a skin wound model

To investigate the antibacterial activity of intracellular Cas12a2, we evaluated an inducible expression system carrying a crRNA directed against the chloramphenicol-resistance transcript in *E. coli* BL21(DE3) pLysS. The active targeting construct (Cas12a2-Chl) was compared with a designated inactive ΔPI-Cas12a2 construct carrying the same targeting crRNA (ΔPI12a2-Chl) and an active Cas12a2 construct without crRNA (Cas12a2-NT). Bacterial growth was monitored under uninduced and induced conditions, and the time required to reach an OD600 of 0.15 was used to quantify growth delay. For Cas12a2-Chl, induction increased the time to threshold from 180 to 441 min, representing a delay of 261 min. In contrast, the corresponding times changed from 123 to 132 min for ΔPI12a2-Chl and from 162 to 183 min for Cas12a2-NT, representing delays of 9 and 21 min, respectively (**Figure 2a**). Thus, induction produced a substantially greater delay in reaching the growth threshold in Cas12a2-Chl than in either control, consistent with a growth-delaying effect associated with functional Cas12a2 and guide-directed recognition of the chloramphenicol-resistance transcript.

**Figure 2.**
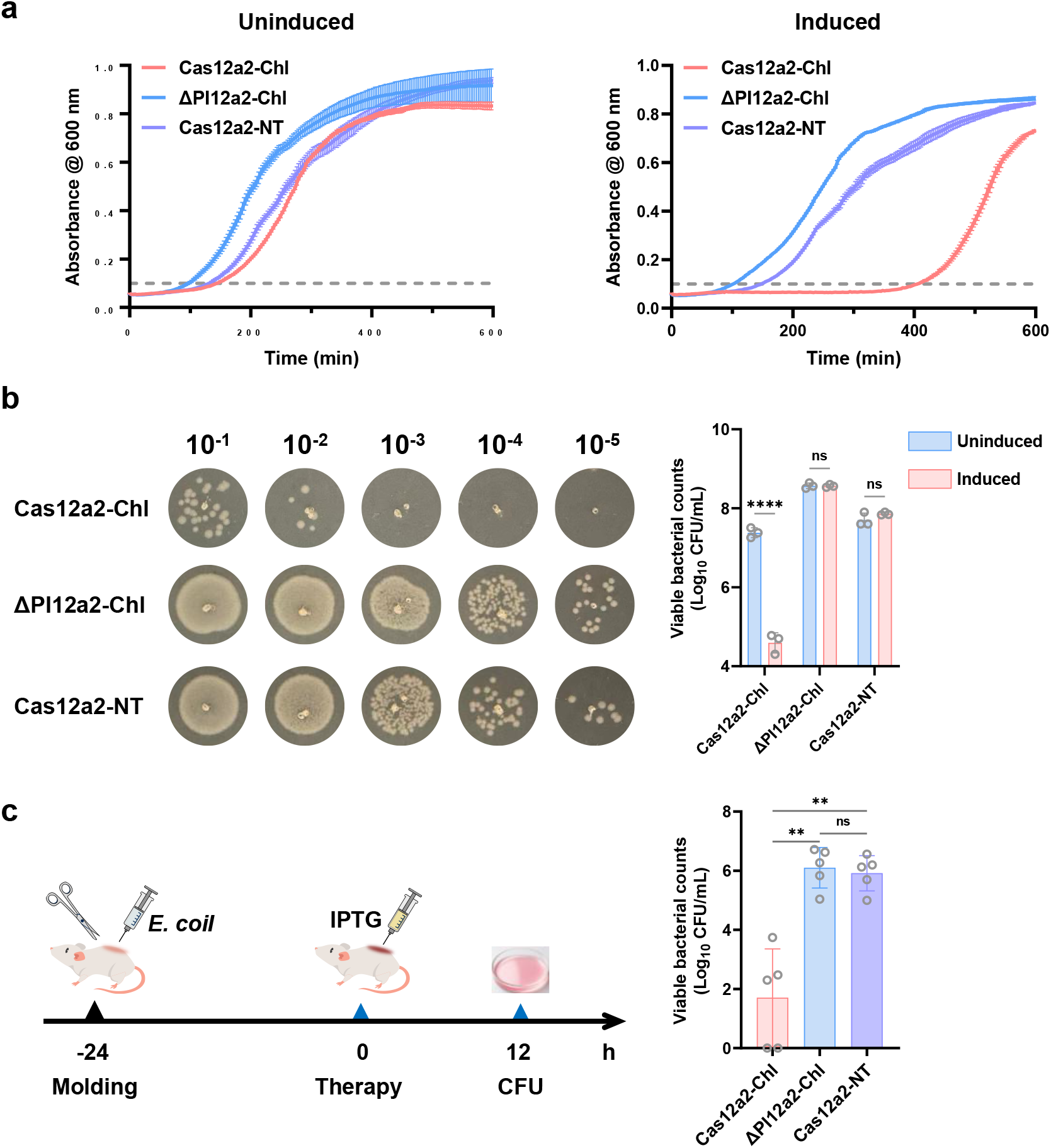
Intracellular Cas12a2 targeting affects bacterial growth and viability in culture and bacterial burden in a skin wound model. (a) Growth curves of bacteria carrying Cas12a2-Chl, ΔPI12a2-Chl, or the non-targeting construct Cas12a2-NT under uninduced and 0.1 mM IPTG-induced conditions (*n* = 3). Growth delay was assessed using the time required to reach an OD600 of 0.15. (b) Representative colony images and quantification of recoverable colony-forming units (CFU) under uninduced and induced conditions (0.1 mM IPTG) (*n* = 3). Log-transformed CFU data were analyzed using two-way ANOVA followed by Šídák’s multiple comparisons test to compare induced and uninduced conditions within each construct. \*\*\*\**P* < 0.0001. (c) Schematic of the mouse skin wound infection model and local IPTG treatment workflow, with quantification of bacterial burdens following treatment (*n* = 5). Wounds were inoculated with bacteria carrying the indicated constructs before IPTG administration. Data were transformed as log10(CFU + 1) and analyzed using one-way ANOVA followed by Dunnett’s T3 multiple comparisons test. \*\*\*\**P* < 0.0001, \*\**P* < 0.01. No colonies were recovered from two of five mice in the Cas12a2-Chl group under the assay conditions; these observations do not establish complete bacterial clearance.

To complement the growth-curve measurements, recoverable viable bacteria were quantified by colony-forming unit (CFU) assays under uninduced and induced conditions. Two-way ANOVA of log-transformed CFU data revealed a significant interaction between construct identity and induction (P < 0.0001), indicating that the effect of IPTG induction differed among the three construct-bearing strains. Šídák’s multiple comparisons test showed that induction with 0.1 mM IPTG reduced recoverable CFU in Cas12a2-Chl by more than 3 log10 units relative to its uninduced control (P < 0.0001). In contrast, no statistically significant induction-associated differences were detected for ΔPI12a2-Chl (P > 0.99) or Cas12a2-NT (P = 0.48) (**Figure 2b**). Together with the pronounced delay in reaching the OD600 threshold, these findings support an antibacterial effect associated with functional Cas12a2 and guide-directed recognition of the chloramphenicol-resistance transcript.

To extend the in vitro findings, we evaluated the pre-installed Cas12a2 system in a mouse skin wound model. Wounds were inoculated with bacteria carrying the indicated constructs, and bacterial burdens were quantified following local IPTG treatment (**Figure 2c**). The Cas12a2-Chl group showed significantly lower bacterial burdens than both the ΔPI12a2-Chl and Cas12a2-NT groups (P = 0.0073, 0.0081 respectively). No significant difference was detected between the two control groups (P = 0.96). These findings support the antibacterial activity of the pre-installed Cas12a2 targeting system in the wound environment.

### Cas12a2 activation is associated with SOS-related transcriptional changes

To identify potential secondary targets for a dual-targeting strategy, we performed RNA sequencing of Cas12a2-Chl and ΔPI12a2-Chl following IPTG induction. Transcriptome comparison revealed a Pearson correlation coefficient of 0.7883, with 272 genes upregulated and 816 genes downregulated in Cas12a2-Chl relative to ΔPI12a2-Chl (**Figure 3a**). These results indicate broad transcriptional differences between bacteria carrying the functional Cas12a2 system and the ΔPI mutant under inducing conditions. We next examined the expression of selected genes associated with the SOS response, DNA repair, and replication, including *recA, ruvA, priB, recR*, and *dnaE*. Among these genes, *recA* and *priB* were significantly upregulated in Cas12a2-Chl compared with ΔPI12a2-Chl (**Figure 3b**). Given the central role of RecA in homologous recombination and SOS response initiation, *recA* was selected as a candidate secondary target.

**Figure 3.**
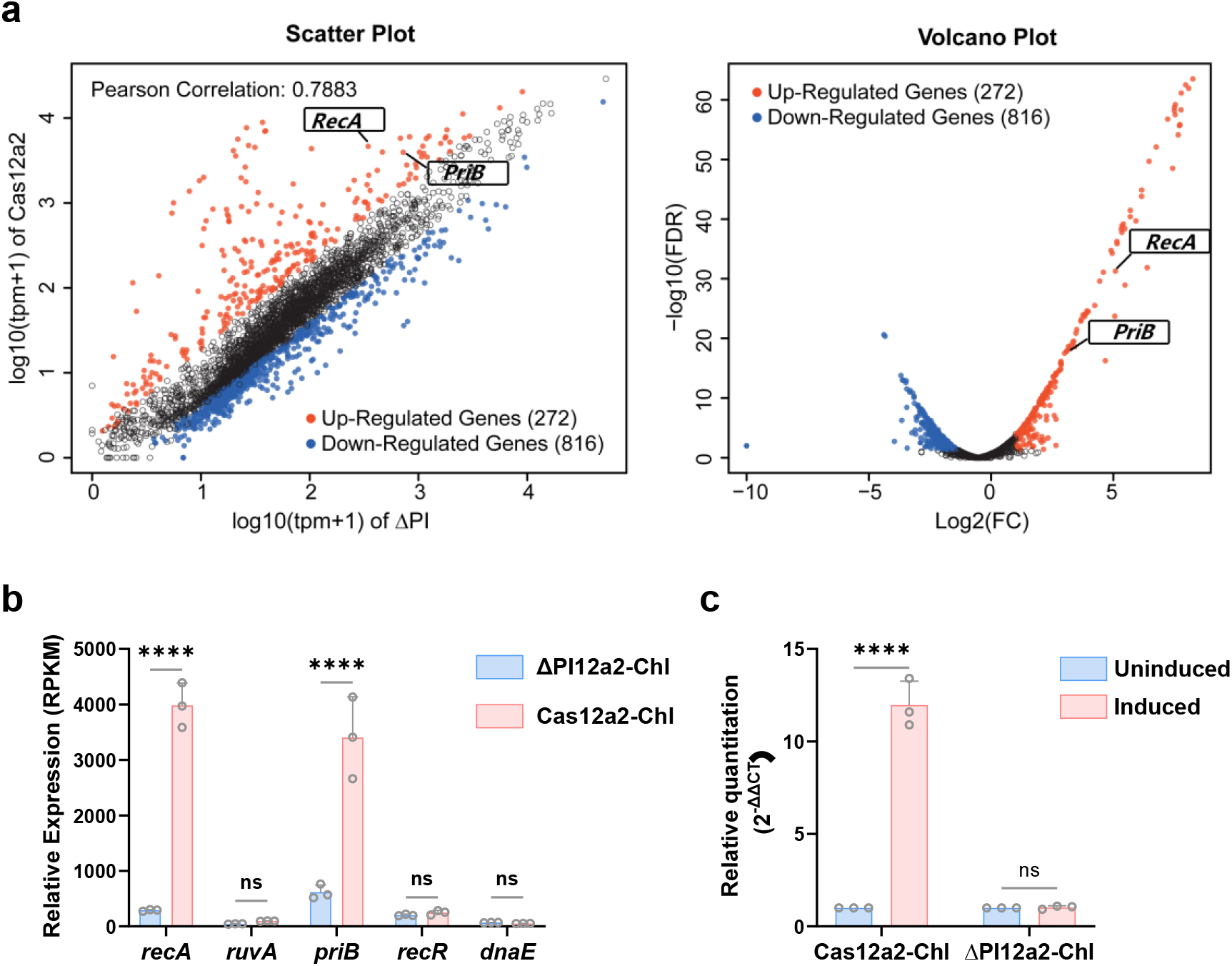
Transcriptional profiling and RT-qPCR validation identify *recA* as a candidate secondary target following Cas12a2 activation. (a) Scatter plot and volcano plot comparing gene expression between Cas12a2-Chl and ΔPI12a2-Chl following IPTG induction. The Pearson correlation coefficient was 0.7883; 272 genes were upregulated and 816 genes were downregulated in Cas12a2-Chl relative to ΔPI12a2-Chl. (b) Expression of selected genes associated with the SOS response, DNA repair, and replication from the RNA-seq dataset. Both *recA* and *priB* were significantly upregulated in Cas12a2-Chl compared with ΔPI12a2-Chl. (c) RT-qPCR validation of *recA* expression in Cas12a2-Chl and ΔPI12a2-Chl with or without IPTG induction. Relative expression was determined using the ΔΔCt method. Data were analyzed using two-way ANOVA followed by Šídák’s multiple comparisons test (*n*=3). \*\*\*\**P* < 0.0001.

To validate the induction-associated change in *recA* expression, we performed RT-qPCR in both strains under induced and uninduced conditions. Relative expression analysis using the ΔΔCt method showed that IPTG induction significantly increased *recA* expression in Cas12a2-Chl, whereas no significant induction-associated change was detected in ΔPI12a2-Chl (**Figure 3c**). Together, these findings link functional Cas12a2 activity to increased *recA* expression and provide a rationale for evaluating *recA* co-targeting in subsequent experiments.

### Dual-targeting of Chl and *recA* enhances Cas12a2-mediated antibacterial activity

To evaluate whether *recA* co-targeting enhances Cas12a2-mediated growth inhibition, we constructed a dual-targeting system directed against both the Chl target and recA. Growth-curve monitoring showed that dual-targeting delayed the time to reach the defined growth threshold by 420 min, compared with 280 min for Chl targeting alone and 15 min for recA targeting alone (**Figure. 4a**). These results demonstrate greater growth inhibition by dual-targeting than by either single-targeting system. It should be noted that single targeting *recA* almost does not cause growth inhibition, which may be due to the insufficient number of *recA* transcripts under normal conditions to trigger Cas12a2 activation.

**Figure 4.**
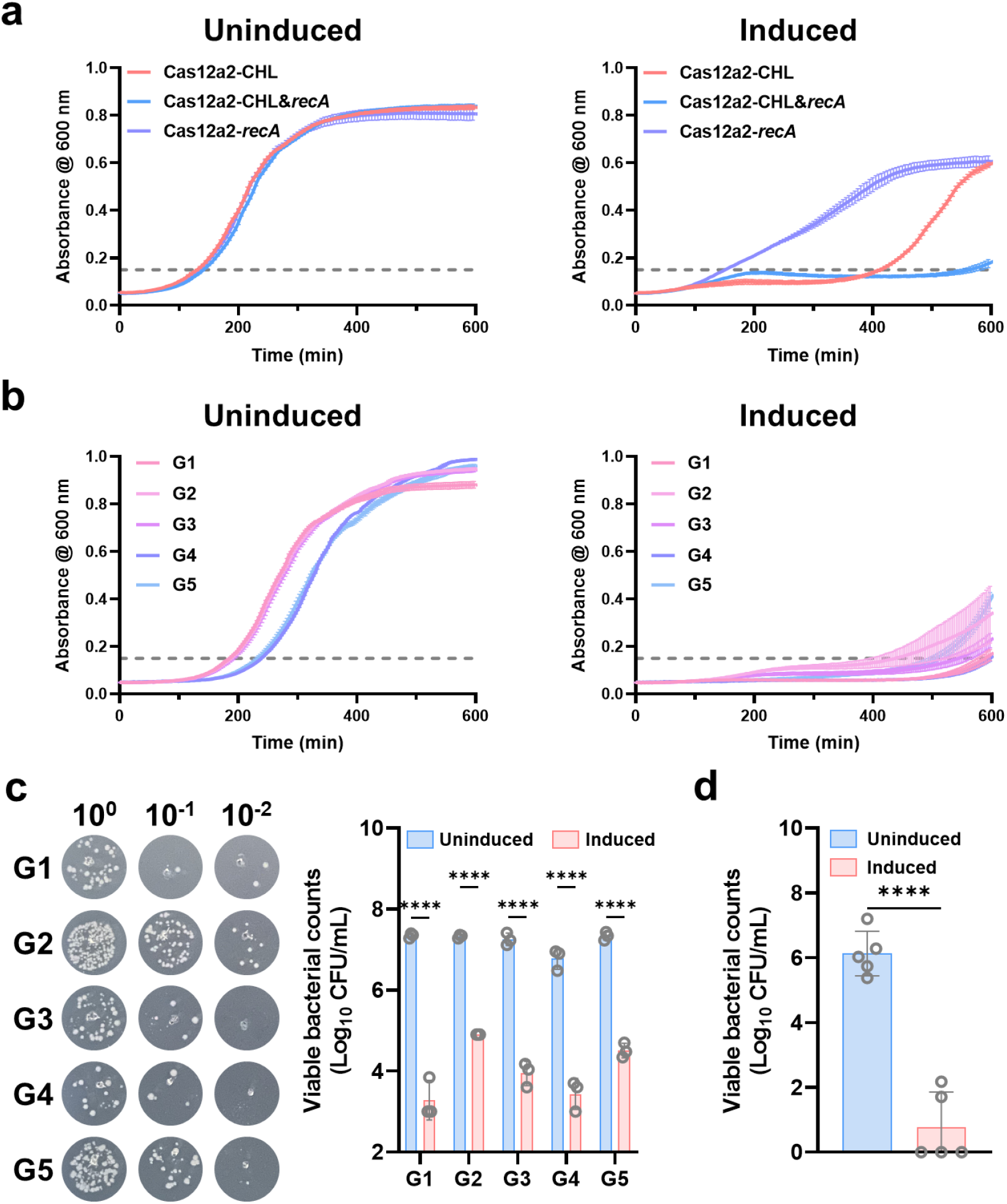
Evaluation and optimization of the antibacterial activity of the Chl-*recA* dual-targeting system. (a) Growth curves comparing Chl single-targeting, Chl-recA dual-targeting, and recA single-targeting systems (*n* = 3). Growth delay was assessed using the time required to reach an OD600 of 0.15. (b) Growth curves of five dual-targeting constructs differing in crRNA sequence composition and number, used to screen for enhanced growth inhibition (*n* = 3). G1: Chl1+RecA1, G2: Chl1*2+RecA1*2, G3: Chl1*3+RecA1*3, G4: Chl1&2+RecA1&2, G5: Chl1&2&3+RecA1&2&3. Growth delay was assessed using the time required to reach an OD600 of 0.15. (c) CFU enumeration for the five dual-targeting constructs (*n*=3). Log-transformed CFU data were analyzed using two-way ANOVA followed by Šídák’s multiple comparisons test to compare induced and uninduced conditions within each construct. \*\*\*\**P* < 0.0001. Growth-curve and CFU analyses were used together to select the preferred crRNA combination for in vivo evaluation. (d) Bacterial burdens determined by CFU enumeration in a mouse skin wound infection model using the selected dual-targeting construct (*n*=5). Statistical significance was determined using an unpaired Welch’s t-test (to account for unequal variances) and further validated by a non-parametric Mann-Whitney *U* test. \*\*\*\**P* < 0.0001.

We next designed three crRNAs for each target and generated dual-targeting constructs with different crRNA combinations and numbers. Screening by growth-curve monitoring and CFU enumeration identified a preferred combination (G1 in Figure 4) based on growth inhibition and reduced recoverable CFU (**Figure. 4b, c**). Finally, we assessed the optimized dual-targeting system in a mouse skin wound infection model. Following IPTG treatment, bacterial burdens were approximately four orders of magnitude lower than those in the untreated group (**Figure. 4d**). Together, these findings support Chl-*recA* dual-targeting as a strategy to enhance Cas12a2-mediated antibacterial activity and demonstrate its efficacy in the wound infection setting.

### Preliminary assessment of resistance-plasmid-associated DNA during SuCas12a2-mediated antibacterial treatment

To investigate whether SuCas12a2-mediated antibacterial activity is accompanied by a reduction in extracellular resistance-plasmid-associated DNA, we designed five qPCR amplicons targeting distinct regions of the resistance plasmid. Culture supernatants were collected from the SuCas12a2-treated, antibiotic-treated, and untreated control groups at 0, 3, and 5 h and analyzed by qPCR (**Figure 5**). At 3 h, reciprocal Ct values increased relative to baseline in all three groups, consistent with an initial accumulation of amplifiable plasmid-associated DNA in the culture supernatants. By 5 h, reciprocal Ct values decreased relative to those at 3 h in the SuCas12a2-treated group, whereas they continued to increase in the antibiotic-treated and untreated groups. This divergent temporal pattern suggests that SuCas12a2-mediated antibacterial treatment was accompanied by a subsequent reduction in extracellular amplifiable plasmid-associated DNA. These findings are consistent with potential degradation of resistance-plasmid DNA during treatment.

**Figure 5.**
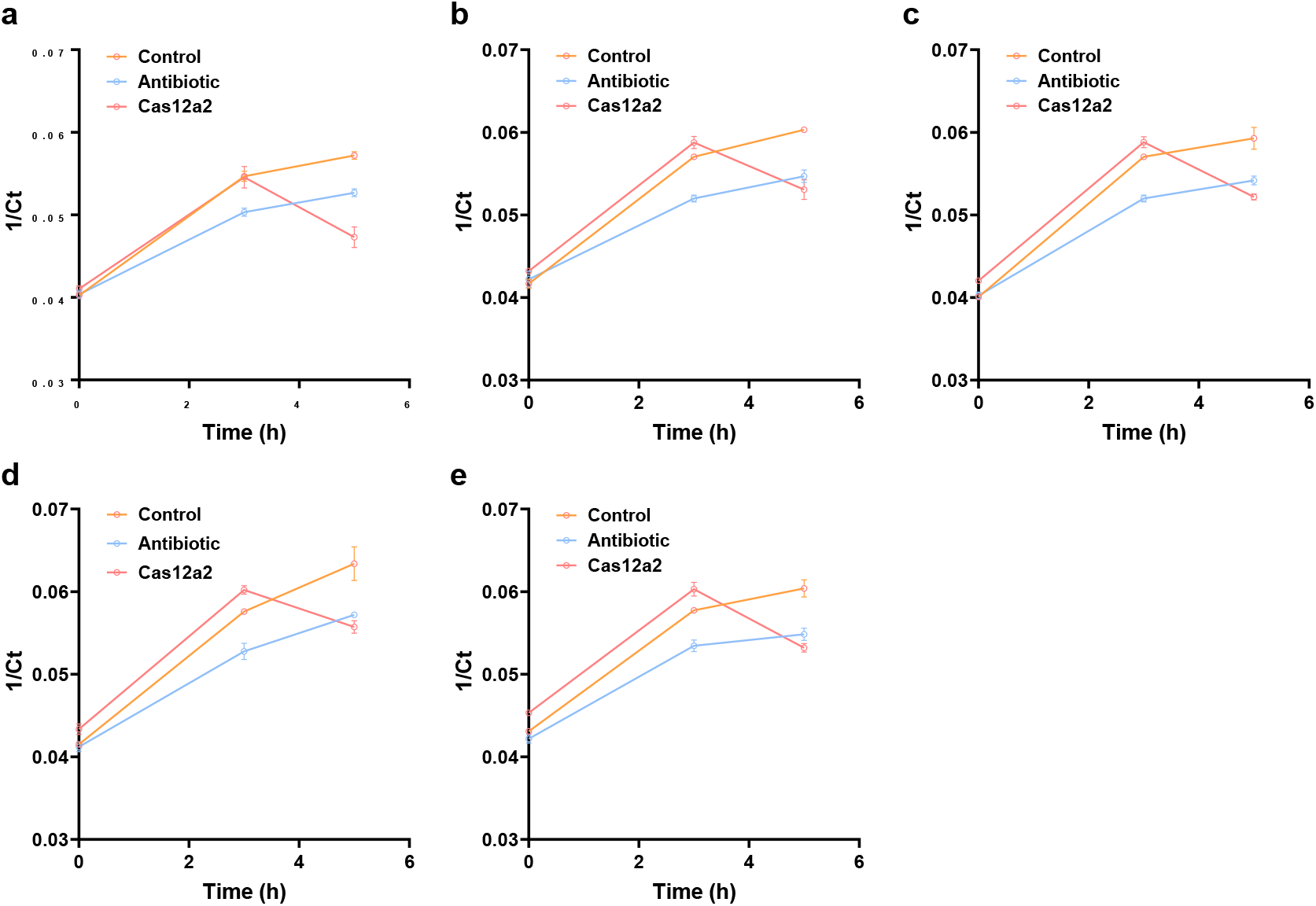
Temporal changes in extracellular resistance-plasmid-associated DNA during SuCas12a2-mediated antibacterial treatment. **(a–e)** qPCR analyses of five distinct regions of the resistance plasmid in culture supernatants collected from the SuCas12a2-treated, antibiotic-treated, and untreated control groups at 0, 3, and 5 h. Each panel represents one amplicon. Results are presented as reciprocal Ct values (1/Ct) (*n*=3).

## Conclusion

In summary, we demonstrated that CRISPR/Cas12a2 programmed against a resistance transcript provides effective, targeted antibacterial activity *in vitro* and *in vivo*. Activation of Cas12a2 induces the SOS response, enabling the identification of *recA* as a secondary target. Co-targeting the resistance transcript and *recA* synergistically enhances bacterial killing. This dual-targeting strategy represents a promising programmable platform to combat AMR and potentially mitigate the spread of resistance plasmids.

## Materials and Methods

### Bacterial strains and expression constructs

*E. coli* BL21(DE3) pLysS (Beyotime), carrying a chloramphenicol-resistance determinant on the pLysS plasmid, was used as the bacterial model. Plasmids encoding suCas12a2 and the indicated crRNA configurations were constructed by GenScript. Expression of Cas12a2/the relevant expression cassette was controlled by an IPTG-inducible T7 promoter system. Two control constructs were evaluated: a ΔPI-Cas12a2 construct carrying the resistance-transcript-targeting crRNA and an active Cas12a2 construct without crRNA. The ΔPI-Cas12a2 construct contained Cas12a2 with the deletion of PFS interacting domain and was designated an inactive control. The non-targeting crRNA contained a random sequence, with its targeting status assessed against based on nucleotide blast.

### Intracellular antibacterial activity assays

The expression constructs were introduced into *E. coli* BL21(DE3) pLysS, and construct-bearing cultures were established using resistance screening. Bacteria were cultured in LB medium under 37°C. The principal comparison included the active resistance-transcript-targeting construct, the ΔPI-Cas12a2 control and the non-targeting crRNA control. Expression was induced with IPTG under 37°C in LB medium. Corresponding cultures without IPTG were evaluated in parallel. Cultures were standardized before comparison by OD600 measurement.

### Bacterial growth measurements

Bacterial growth was monitored by measuring OD600 using a Multiskan FC microplate photometer (Thermo Fisher Scientific). Measurements were collected at 3 min over 10 h under 37°C in LB medium. Each biological replicate was an independently established bacterial culture.

### Viable bacterial counts

Samples were collected at 10 h for culture-based enumeration. Bacteria were recovered on agar medium, with the presence or absence of selective agents specified for each experiment. Colony counts were converted to CFU/mL using the plated volume and dilution factor. Their treatment in graphical displays and statistical analyses is described in the Statistical analysis section. Antibacterial effects were expressed as absolute CFU values differences relative to a specified comparator.

### RNA sampling and transcriptome sequencing

For transcriptome analysis, Cas12a2-Chl and ΔPICas-Chl cultures were sampled at 5 h after IPTG induction, with three independent biological replicates per group. Total RNA was extracted from bacterial samples using TRIzol reagent (Invitrogen). RNA quantity and quality were assessed using a NanoDrop ND-2000 spectrophotometer and an Agilent 2100 Bioanalyzer. Following rRNA depletion, strand-specific libraries were prepared using TruSeq RNA sample preparation Kit from Illumina (San Diego, CA). Libraries were sequenced on an Illumina NovaSeq 6000 platform to generate 150-bp paired-end reads.

Differential expression analysis was performed using edgeR with gene-level count data. Differentially expressed genes were defined using an absolute log_2_ fold-change threshold of 2 and a false discovery rate below 0.05. Normalized expression values used for visualization were calculated separately from the count-based differential expression analysis. GO and KEGG enrichment analyses were conducted using Goatools and KOBAS, respectively.

### RT-qPCR analysis of *recA* expression

Relative *recA* transcript abundance was assessed by RT-qPCR. RNA preparation and DNA removal were performed as described above. Complementary DNA was synthesis using HiScript II qRT SuperMixII (Vazyme) under 50 °C for 15 min, and quantitative PCR was conducted using on ABI 7300 plus (Thermo Fisher Scientific). Relative expression was calculated using ΔΔCt method. Each biological replicate comprised an independent bacterial culture. RT-qPCR values were interpreted as relative transcript abundance and not as absolute transcript numbers per bacterium.

### Mouse skin wound model

Animal procedures were approved by Ethics Committee of China Pharmaceutical University and performed in accordance with guidelines. Mice were 8-week-old BALB/c mice weighing 18-20 g. Anaesthesia, analgesia, monitoring, humane endpoints and euthanasia procedures were conducted according to the approved protocol. The wound model was established using a biopsy punch (typically 5 mm in diameter). Wounds received *E. coli* carrying the relevant expression constructs before inoculation. Groups comprised bacteria carrying the active resistance-transcript-targeting construct, the ΔPI-Cas12a2 control or the non-targeting crRNA control. Local IPTG administration was used to induce the pre-installed expression system. Wound samples were collected at 12 h after IPTG treatment. Recovered bacteria were enumerated on MacConkey agar.

## Acknowledgments

This work was supported by Special Fund of State Key Laboratory of Respiratory Health and Multimorbidity (2060204); National Natural Science Foundation of China (61901527); National Innovation and Entrepreneurship Training Program for Undergraduate; the Beijing Association of Holistic Integrative Medicine Project (ZHKY-2026-112 (B006)).

